# Measures of neural similarity

**DOI:** 10.1101/439893

**Authors:** S. Bobadilla-Suarez, C. Ahlheim, A. Mehrotra, A. Panos, B. C. Love

## Abstract

One fundamental question is what makes two brain states similar. For example, what makes the activity in visual cortex elicited from viewing a robin similar to a sparrow? One common assumption in fMRI analysis is that neural similarity is described by Pearson correlation. However, there are a host of other possibilities, including Minkowski and Mahalanobis measures, with each differing in its mathematical, theoretical, neural computational assumptions. Moreover, the operable measures may vary across brain regions and tasks. Here, we evaluated which of several competing similarity measures best captured neural similarity. Our technique uses a decoding approach to assess the information present in a brain region and the similarity measures that best correspond to the classifier’s confusion matrix are preferred. Across two published fMRI datasets, we found the preferred neural similarity measures were common across brain regions, but differed across tasks. Moreover, Pearson correlation was consistently surpassed by alternatives.

## 1. Introduction

Detecting similarities is critical to a range of cognitive processes and tasks, such as memory retrieval, analogy, decision making, categorization, object recognition, and reasoning [1, 2, 3, 4, 5, 6]. Key questions for neuroscience include which measures of similarity does the brain use, and do similarity computations differ across brain regions and tasks. Whereas psychology has considered a dizzying array of competing accounts of similarity [7, 8, 9, 10, 11, 12, 13], research in neuroscience usually assumes that Pearson correlation captures the similarity between different brain states [14, 15, 16, 17, 18, 19, 20, 21]), though see [22, 23, 24, 16].

On the face of it, it seems unlikely that the brain would use a single measure of similarity across regions and tasks. First, across regions, the signal and type of information represented can differ [6, 25, 26], which might lead the accompanying similarity operations to also differ. Second, task differences, such as those that shift attention [27, 28, 29], lead to changes in the brain’s similarity space which may reflect basic changes in the underlying similarity computation. Outside neuroscience it is common to use different similarity measures on different representations. For example, in machine learning, Euclidean measures are often used to determine neighbors in image embeddings whereas cosine similarity is more commonly used in natural language processing [30].

In this contribution, we developed a technique to address two specific goals. The first goal was to ascertain whether the similarity measures used by the brain differ across regions. The second goal was to investigate whether the preferred measures differ across tasks and stimulus conditions. Our broader aim was to elucidate the nature of neural similarity.

Previous studies have adopted different similarity measures to relate pairs of brain states such as Pearson correlation or the Mahalanobis measure [31, 32, 33, 14]. However, the basis for choosing one measure over another is not always clear. The choice of measure induces a host of assumptions, including assumptions about how the brain codes and processes information. While all the measures considered operate on two vectors associated with two brain states (e.g., the BOLD response elicited across voxels when a subject views a truck vs. a moped), the operations performed when comparing these two vectors differ for each similarity measure.

### 1.1. Families of similarity measures

To better understand these assumptions and their importance, we organise common measures of similarity, many of which are used in the neuroscience literature, into three families (see Figure 1, left side). The most basic split is between similarity measures that focus on the angle between vectors (e.g., Pearson correlation or cosine distance) and measures that focus on differences in vector magnitudes. The latter branch subdivides between distributional measures that are sensitive to covariance across vector dimensions (e.g., Mahalanobis) and those that are not (e.g., Euclidean). Of course there are uncountably infinite similarity measures one could choose to assess; the goal here is to compare common measures that can discriminate between different computations of interest as organized by these families of measures with focus on angle, magnitude, and distributional properties.

**Figure 1:**
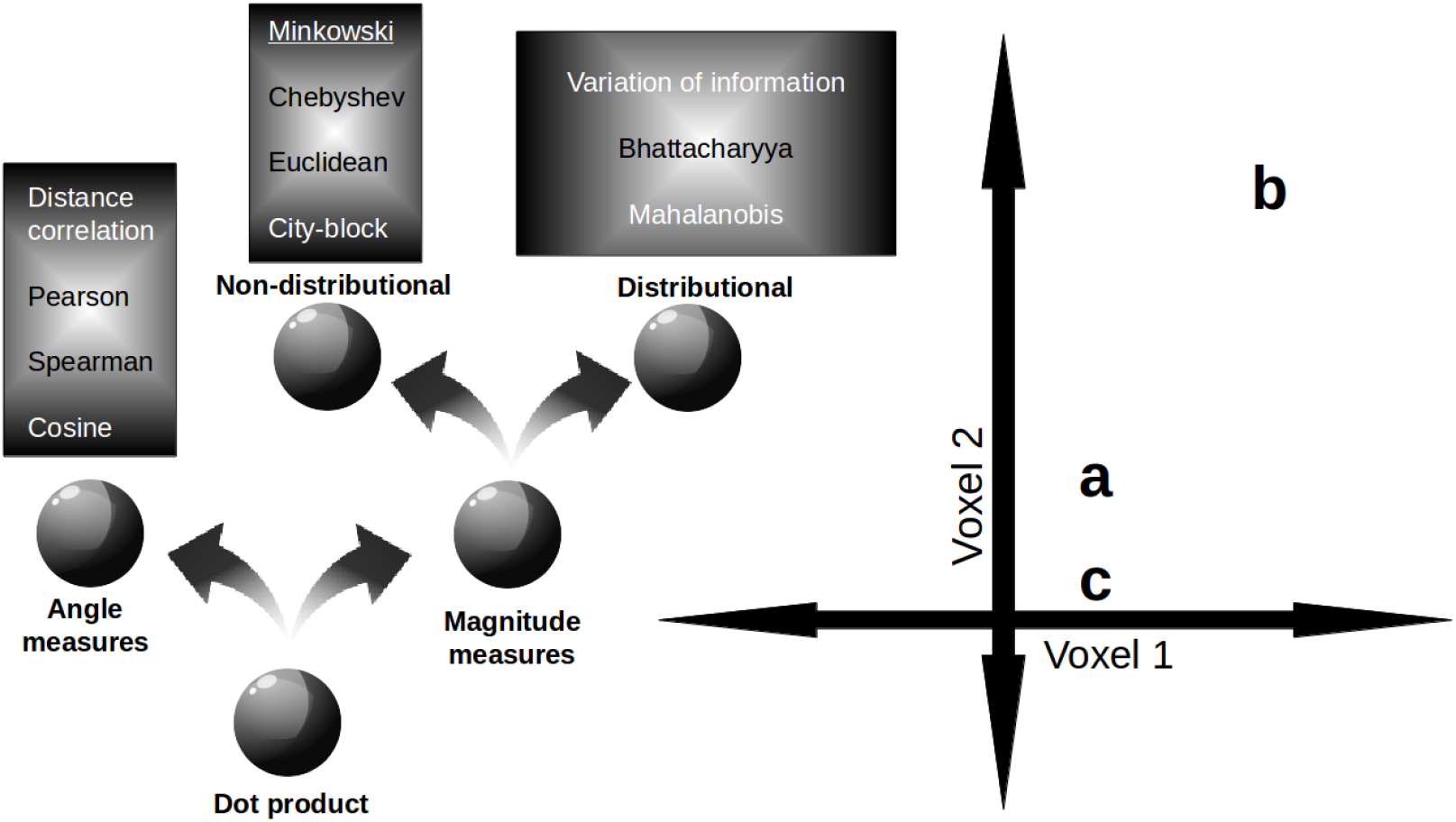
Families of similarity measures. (left panel) Similarity measures divide into those concerned with angle vs. magnitude differences between vectors. Pearson correlation and Euclidean distance are common angle and magnitude measures, respectively. The magnitude family further subdivides according to distributional assumptions. Measures like Mahalanobis are distributional in that they are sensitive to co-variance such that similarity falls more rapidly along low variance directions. (right panel) The choice of similarity measure can strongly affect inferences about neural representational spaces. In this example, stimuli a, b, and c elicit different patterns of activity across two voxels. When Pearson correlation is used, stimulus a is more similar to b than to c. However, when the Euclidean measure is used, the pattern reverses such that stimulus a is more similar to c than b.

The choice of similarity measure can shape how neural data are interpreted. Consider the right panel in Figure 1. In this example, the neural representation of object a is more similar to that of b than c when an angle measure is used, but this pattern reverses when a magnitude measure is used.

Unlike the other measures, distributional measures are anisotropic, meaning the direction of measurement is consequential.^1^ Examples of such measures are variation of information, Mahalanobis, and Bhattacharyya measures. These measures consider the covariance between stimuli dimensions, which implies that the direction (in feature or voxel space) along which the measurement is made will impact the measurement itself.

The choice of similarity measure reflects basic assumptions about the nature of the underlying neural computation. For example, Pearson correlation (a common measure for neural similarity in fMRI, e.g., [14, 15, 16, 17, 18, 19, 20, 35]) assumes that overall levels of voxel activity are normalized and that each voxel independently contributes to similarity, whereas Minkowski measures assume similarity involves distances in a metrical space instead of vector directions. Furthermore, the Mahalanobis measure expands on both Minkowski and Pearson by assuming that the distributional pattern of voxel activity is consequential.

Knowing which similarity measure best describes the brain’s operation would not only improve data analyses, but could also illuminate the nature of neural computation at multiple levels of analysis. For example, if a brain region normalized input patterns for key computations, then Pearson correlation might have superior descriptive power than the dot product. At a lower level, such a result would be consistent with mutually inhibiting single cells [36]. On the other hand, if the brain matches to a rigid template or filter (e.g., [37]), then the Euclidean measure should provide a better explanation for neural data.

To identify which similarity measures are used by the brain requires addressing a number of challenges. One challenge is to specify a standard by which to evaluate competing similarity measures. Related work in Psychology and Neuroscience has relied on evaluating against verbal report. How ever, such an approach is not suited to our aims because we are interested in neural computations that may differ across brain regions and which may not be accessible by verbal report or introspection.

Instead, we rely on a decoding approach to assess the information latent in a brain region. The intuition is that brain states that are similar should be confusable in decoding. For example, a machine classifier may be more likely to confuse the brain activity elicited by a bicycle with that by a motorcycle than a car. In this fashion, we can evaluate competing similarity measures on a per region basis in a manner that is not constrained by verbal report. The insight that similarity is intimately related to confusability has a long and rich intellectual history [38, 39, 40] though has not yet been considered to evaluate what makes two brain states similar.

### 1.2. Discrimination of similarity measures

Our method for distinguishing the similarity measure used by the brain involves two basic steps:

1. For each ROI, compute a pairwise confusion matrix using a classifier. For each ROI, also compute a similarity matrix for each candidate similarity measure.
2. For each similarity measure, correlate its similarity matrix with the confusion matrix using Spearman correlation to avoid scaling issues.

The better a similarity measures characterizes what makes two brain states similar, the higher its Spearman correlation with the confusion matrix should be. This analysis uses the confusion matrix as an approximation of what information is present in a brain region (more on this below).

The matrices for each similarity measure were optimized to maximize the Spearman correlation with the confusion matrix by performing feature selection on voxels (see Figure 2). See the SI (Supplemental Information) for details on the similarity measures. Importantly, to understand the results, some similarity measures that estimate covariance matrices are tagged according to the type of regularization used; with (d) for keeping only the diagonal entries and (r) for Ledoit-Wolf shrinkage.

**Figure 2:**
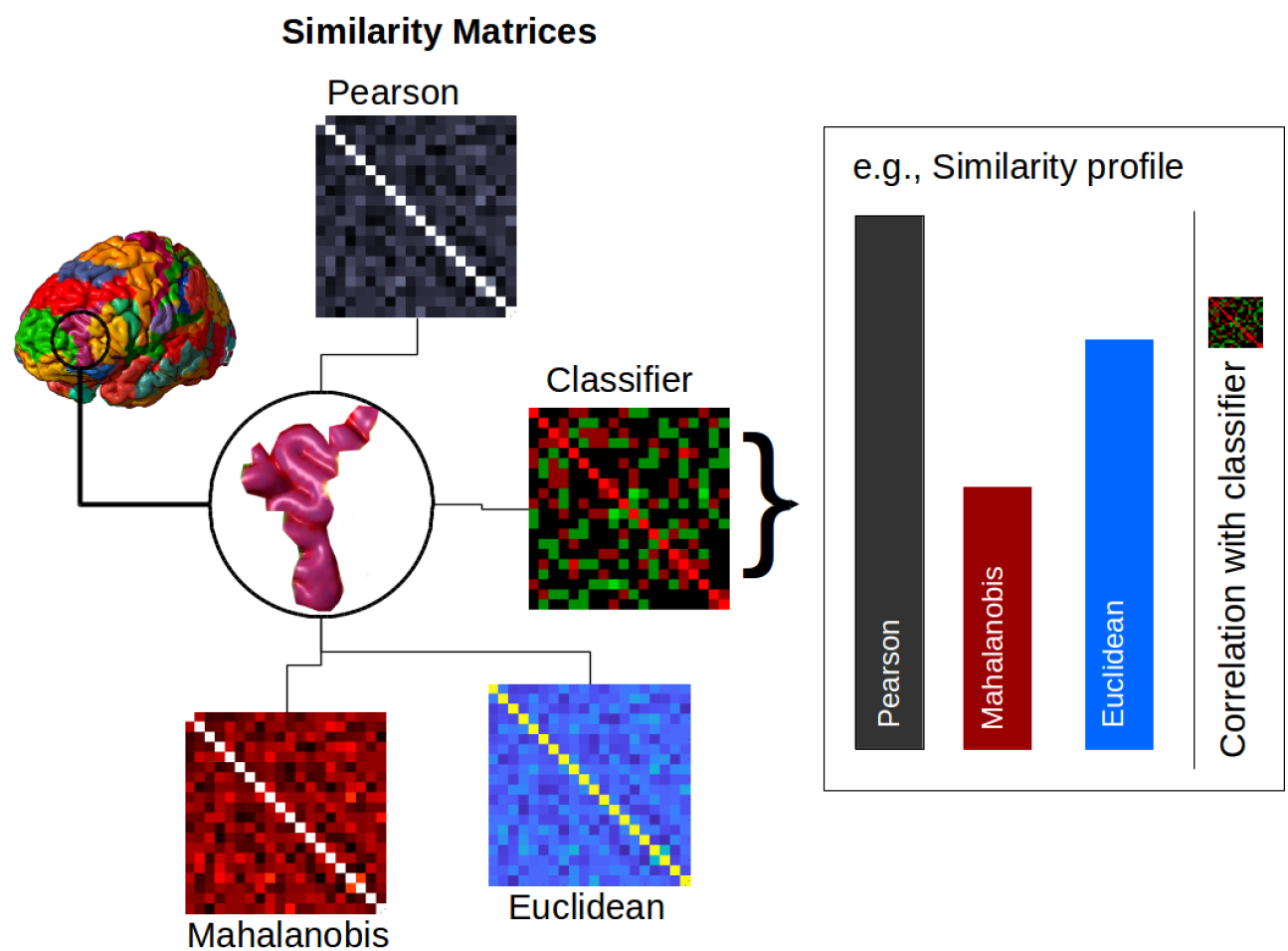
Evaluating the similarity profile for a ROI. The confusion matrix from a classifier is used to approximate the information present in the ROI. The similarity matrix from each similarity measure is correlated with this confusion matrix (i.e., the classifier matrix in the figure). The pattern of these correlations (i.e., the performance of the various similarity measures) is the similarity profile for that ROI. Similarity profiles can be compared between ROIs, both within and between datasets (see Materials and Methods section for more details).

We considered all 110 regions of interest (see SI for a list of the 110 regions) from the Oxford-Harvard Brain Atlas (provided with FSL, [41]) for two previously published datasets. One dataset was from a study in which participants viewed geometric shapes (GS) [28] and the other dataset was from a study in which participants viewed natural images (NI) [6]. For each dataset, we determined the top 10 ROIs for decoding accuracy (cf. Bhandari et al. [42]). The union of these top ROIs provided 12 ROIs that were considered in subsequent analyses (see SI).

### 1.3. Lower confusability as information gain

As mentioned above, our proposed method involves approximating brain state information with a classifier. Subsequently, we use this approximation to assess an array of similarity measures. The motivation for using a classifier to approximate information in a brain state arises from an information theoretic perspective. For example, suppose one’s prior assumption is that two stimuli are equally likely, which corresponds to random guessing or maximal entropy (1 bit). If a probabilistic classifier with the same prior is applied to the stimulus and approaches 100% accuracy, then the information gain approaches 1 bit. Formally, one can measure the Kullback-Leibler (KL) divergence (a continuous, non-saturating measure) between a prior distribution p (centered at 0.5) and an updated distribution q defined by the classifier’s output. With a suitable prior distribution for the classifier, the KL-divergence is always defined and enables a computable measure of brain state information. Thus, KL-divergence, or information gain, will be inversely proportional to confusability as measured by the classifier. Of course, in practice, machine classifiers do not reach close to 100% accuracy with fMRI data for the types of discriminations that we consider. The point is that decoding and measuring available information in a brain state are intimately linked.

### 1.4. Classification is not similarity

Although it should be clear cognitive scientists of all varieties that similarity and classification are conceptually distinct (see [2]), it may not be as apparent to some neuroscientists whose focus is elsewhere. To view similarity and classification as one in the same, would be akin to viewing any operation in which similarity could be relevant, such as memory retrieval, as synonymous with similarity [1].

Mathematically, the domain and range of similarity and classification functions are distinct. Similarity takes as its domain (i.e., input) two states and its range (i.e., output) is a scalar value (i.e., the similarity). Notice that similarity can apply to any two states, irrespective of class membership. A similarity function does not need to be “trained” and “tested” on a particular discrimination, but instead can apply broadly. In contrast, a classification function takes as its domain (i.e., input) items drawn from a predetermined set of classes and its range (i.e., output) is a nominal value indicating the class membership of the item. A classifier is trained on items from the contrasting classes and tested only on items drawn from these same distributions.

To showcase the distinction between similarly and classification operators, in addition to our main results, we also present a results for a nonclassification task that relies on neural similarity. In particular, we present results for a triplet task in which we assess whether neural similarity between a standard stimulus and two probe stimuli, one of which matches in shape. The similarity measures that perform best in the triplet task are the ones that perform best in our main decoding analyses. Critically, the stimulus classes used in the triplet task were not included in the decoding analysis, which highlights that similarity functions apply more broadly than classification functions and that our method for selecting the brain’s preferred similarity functions generalizes to novel stimulus classes. Before visiting this result, we present the main results that answer key questions, such as whether the brain’s preferred similarity measures are common across regions and tasks.

## 2. Results

### 2.1. Neural similarity

What makes two brain states similar and does it vary across brain regions and tasks? The following analyses focus both on the performance of individual similarity measures and on the pattern of performance across a set of candidate measures, which we refer to as the *similarity profile* for an ROI (see Figure 2).

As a precursor, we first tested whether similarity measures differed in their performance (Figure 3a). Specifically, we evaluated whether certain measures better describe what makes two brain states similar by nested comparison using a mixed-effects model for each study (see Materials and Methods). For both studies, similarity measures differed in their performance, χ^2^(2) = 1720.331, *p* < 0.001; χ^2^(2) = 6770.249, *p* < 0.001, for the GS and NI studies, respectively.

**Figure 3:**
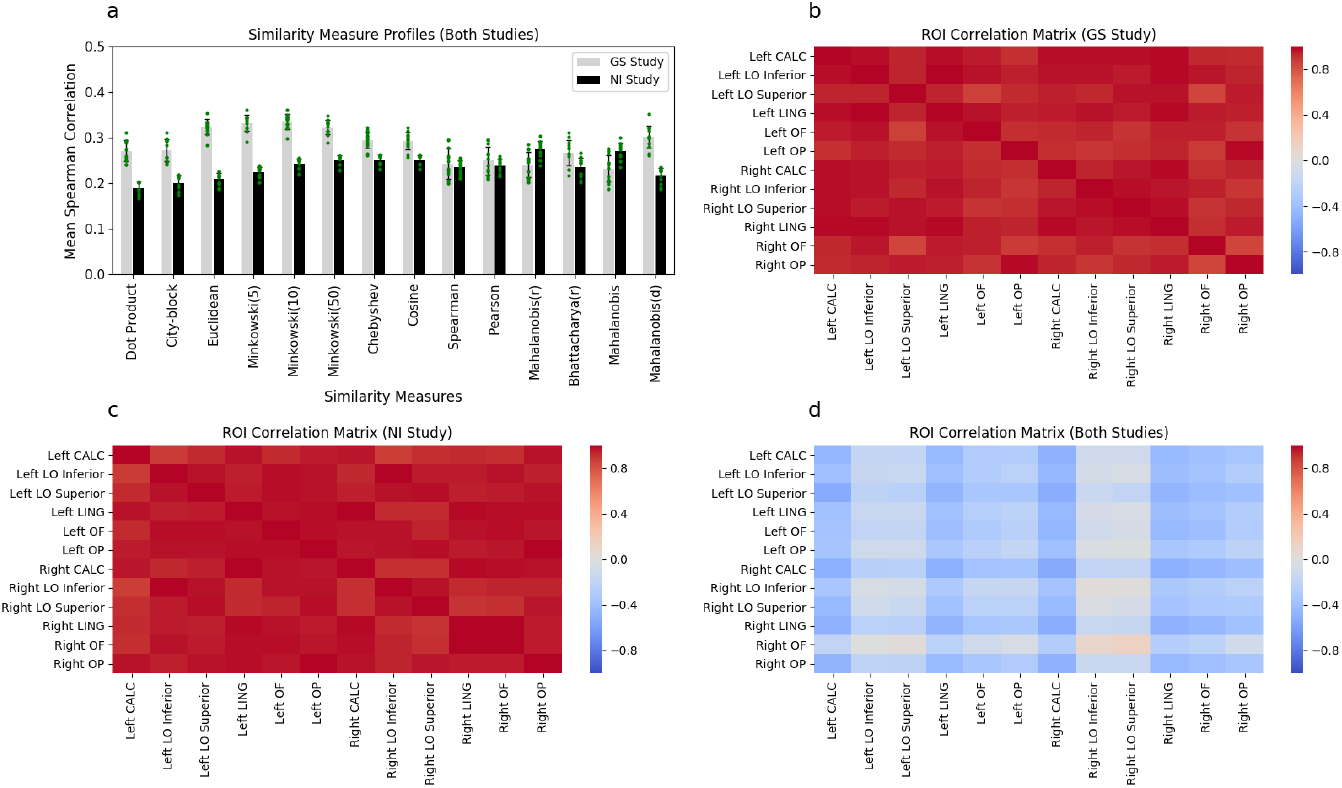
Similarity measure profiles and ROI correlation matrices. Mean Spearman correlations (a) for each similarity measure and the classifier confusion matrix in the GS study (grey bars) and the NI study (black bars) are displayed. To convey the variability, error bars are plotted as standard deviations and each ROI mean is plotted as a green point. ROI correlation matrices for the (b) GS and (c) NI studies, demonstrating that the similarity profiles were alike across brain regions (i.e., were positively Pearson correlated). ROI correlation matrix (d) demonstrating that the similarity profiles disagreed across studies (i.e, were negatively Pearson correlated). The 12 ROIs were left and right intracalcarine cortex (CALC), left and right lateral occipital cortex (LO) inferior and superior divisions, left and right lingual gyrus (LING), left and right occipital fusiform gyrus (OF), and left and right occipital pole (OP).

We tested whether the similarity profile differed across brain regions within each study. The similarity profiles (i.e., mean aggregate performance across measures) were remarkably alike across ROIs (see Materials and Methods). High (Pearson) correlations are presented within task for both the GS study (Figure 3b) and the NI study (Figure 3c) between all pairs of ROIs; where mean correlation of the upper triangle is 0.95 (s.d. = 0.034) in the former and 0.96 (s.d. = 0.027) in the latter. Bartlett’s test [43], which evaluates whether the matrices are different from an identity matrix, was significant for both the GS study, χ^2^(66) = 432.847, *p* < 0.001, and the NI study, χ^2^(66) = 502.7494, *p* < 0.001. Permutation tests (with 10,000 iterations), where the labels of the similarity measures were permuted, confirmed these results (*p* < 0.001). These results are consistent with the same similarity measures being used across brain regions within each study.

We tested whether similarity profiles differed between studies. The results indicated that similarity profiles differed between studies, suggesting that the operable neural similarity measures can change as a function of task or stimuli (Figure 3d). In particular, similarity profiles between studies were negatively correlated with a mean correlation of the upper triangle of −0.27 (s.d. = 0.148). Jennrich’s test [44] showed that this matrix was different than a matrix of zeros, χ^2^(66) = 769.0349, *p* < 0.001. Permutation tests (10,000 iterations) with shuffling of similarity label measures also confirmed these results (*p* < 0.001).

### 2.2. Searchlight analysis

In light of these results, *post hoc* pairwise tests of each similarity against the Pearson similarity measure, which is the *de facto* default choice in the literature, were conducted. The contrasts from the mixed effects models (mentioned above, see Materials and Methods) presented in Table 1 provide evidence that some similarity measures are a superior description of the brain’s similarity measure. The performance of many measures differed from Pearson, especially in the NI study. Notably, only two variants of the Ma-halanobis measure and three Minkowski measures outperformed Pearson. In the GS study, we can observe that all the Minkowski distances performed better than Pearson as well as cosine, Mahalanobis(d), and the dot product. Once again, the contrasting pattern of results between the two studies is striking.

**Table 1:**
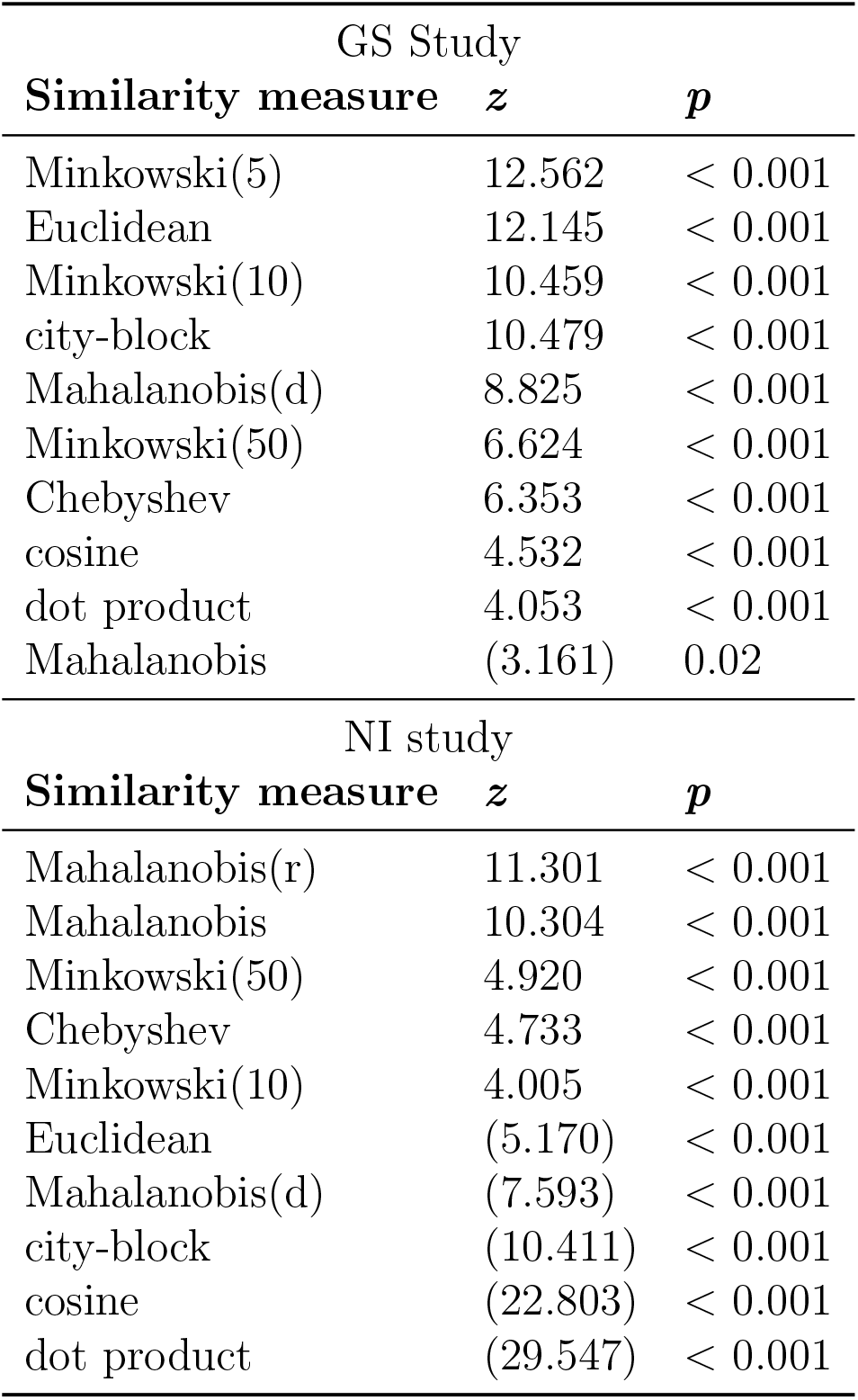
Comparison of similarity measures to Pearson correlation. Top panel shows significant *z* statistics for measures worse than Pearson correlation (in brackets) and better than Pearson correlation for the GS study. Bottom panel shows the same for the NI study. p-values are Bonferroni corrected.

| GS Study |  |  |
| --- | --- | --- |
| Similarity measure | $z$ | $p$ |
| Minkowski(5) | 12.562 | $< 0.001$ |
| Euclidean | 12.145 | $< 0.001$ |
| Minkowski(10) | 10.459 | $< 0.001$ |
| city-block | 10.479 | $< 0.001$ |
| Mahalanobis(d) | 8.825 | $< 0.001$ |
| Minkowski(50) | 6.624 | $< 0.001$ |
| Chebyshev | 6.353 | $< 0.001$ |
| cosine | 4.532 | $< 0.001$ |
| dot product | 4.053 | $< 0.001$ |
| Mahalanobis | (3.161) | 0.02 |
| NI study |  |  |
| Similarity measure | $z$ | $p$ |
| Mahalanobis(r) | 11.301 | $< 0.001$ |
| Mahalanobis | 10.304 | $< 0.001$ |
| Minkowski(50) | 4.920 | $< 0.001$ |
| Chebyshev | 4.733 | $< 0.001$ |
| Minkowski(10) | 4.005 | $< 0.001$ |
| Euclidean | (5.170) | $< 0.001$ |
| Mahalanobis(d) | (7.593) | $< 0.001$ |
| city-block | (10.411) | $< 0.001$ |
| cosine | (22.803) | $< 0.001$ |
| dot product | (29.547) | $< 0.001$ |

Given the performance of the Euclidean and Mahalanobis(r) measures, and that they have been used previously in analyzing neural data [16, 45, 46, 47], we selected these measures for inclusion in a searchlight analysis (Figure 4, see Materials and Methods for details). By comparing the Euclidean and Mahalanobis(r) measures to Pearson correlation on a voxel-by-voxel basis for the 12 ROIs, we aimed to provide a visualization of the performance of similarity measures across regions and studies. Figure 4 illustrates the regions where these two measures outperform Pearson correlation, displaying the maximum *t* for voxels where both Euclidean and Mahalanobis overlap (see SI for visualizations of the overlap).

**Figure 4:**
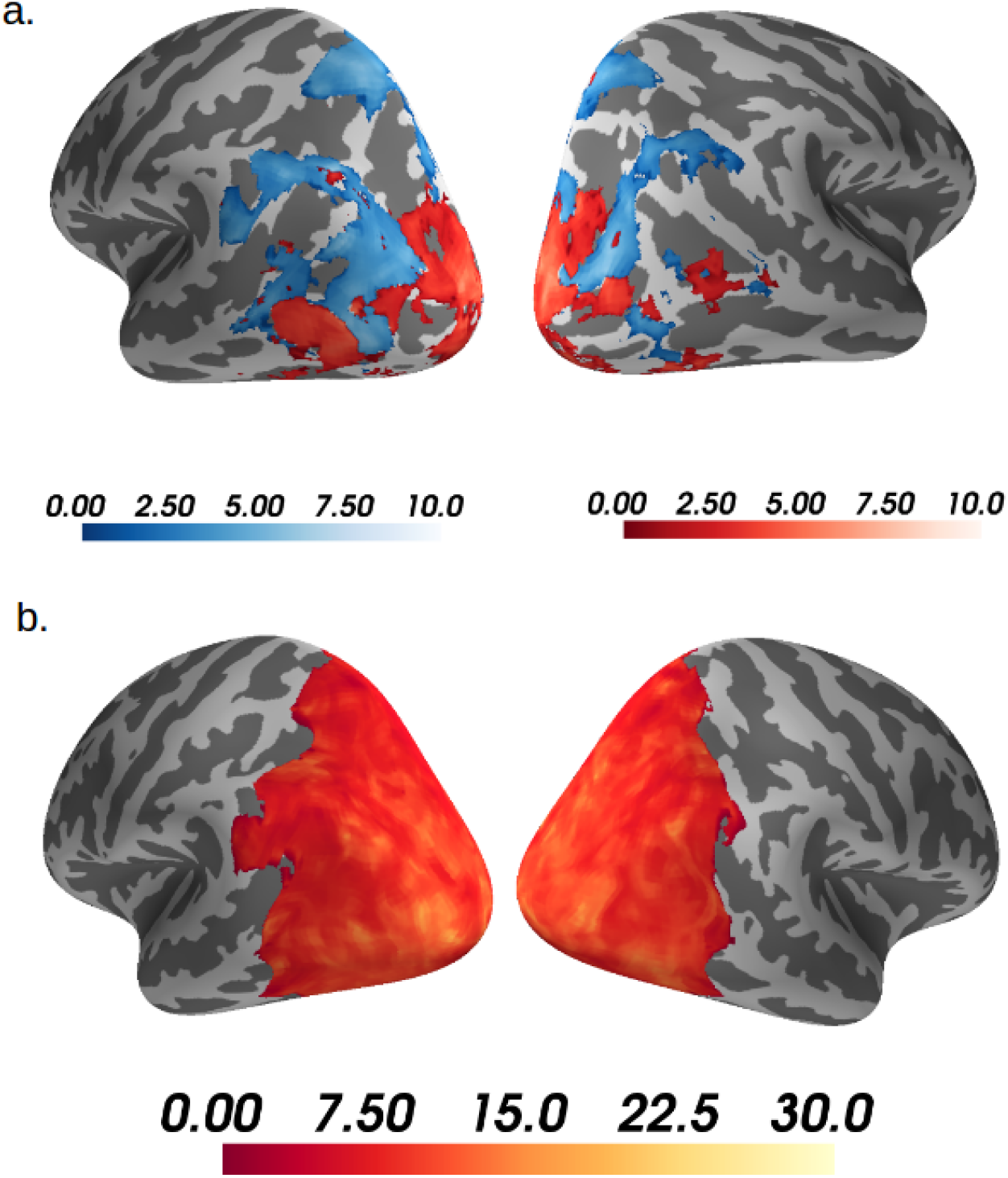
Euclidean & Mahalanobis(r) outperform Pearson. Occipito-lateral views of the left and right hemispheres for the GS study (a) and the NI study (b) displaying maximum t statistics where either the Euclidean measure (blue) or the Mahalanobis(r) measure (red) outperformed the Pearson correlation measure (i.e., each voxel displays the t statistic for the measure with highest t). The t statistics were based on a searchlight analysis of Spearman correlations of each measure with each voxel’s SVM confusion matrix (see Materials and Methods). Only displaying t statistics where *p* < 0.001 for paired sample t-tests, TFCE corrected; computed with FSL’s randomise function with 5000 permutations, using as a mask the 12 ROIs with best accuracy (see Materials and Methods). Note: very few voxels only show the Euclidean measure significantly outperforming Pearson correlation in the NI study, thus do not appear in this visualization.

In the NI study, the Mahalanobis(r) measure dominated (Figure 4b), confirming the results from the previous analyses. In contrast, in the GS study (Figure 4a) Euclidean dominates in some regions whereas Mahalanobis(r) dominates in others. Despite it being a *de facto* standard, Pearson similarity was never the top measure. For this *post hoc* analysis, the measures were compared using permuted paired sample *t* statistics for each voxel. Positive *t* statistics that survived threshold-free cluster enhancement (TFCE) correction with *p* < 0.001 are presented in Figure 4 (see Materials and Methods for the rationale behind this threshold).

### 2.3. Triplet task

As discussed in the Introduction, similarity and classification are distinct concepts. To illustrate, we show how similarity measures can be used in non-classification tasks involving stimuli from novel (untrained) classes. In particular, we consider a triplet task involving data from the NI study (Figure 5). The task is to decide which of two probe items is more neurally similar to the standard stimulus. Trials are defined as correct when the probe that matches in shape is more neurally similar. To foreshadow our results, neural measures that perform best in our decoding analysis perform best in the triplet task, despite the entire classes used in the triplet task being withheld from the decoding analysis. These results indicate that approximating the information available in a brain state through decoding can select similarity measures that broadly generalize and perform sensibly in novel tasks.

**Figure 5:**
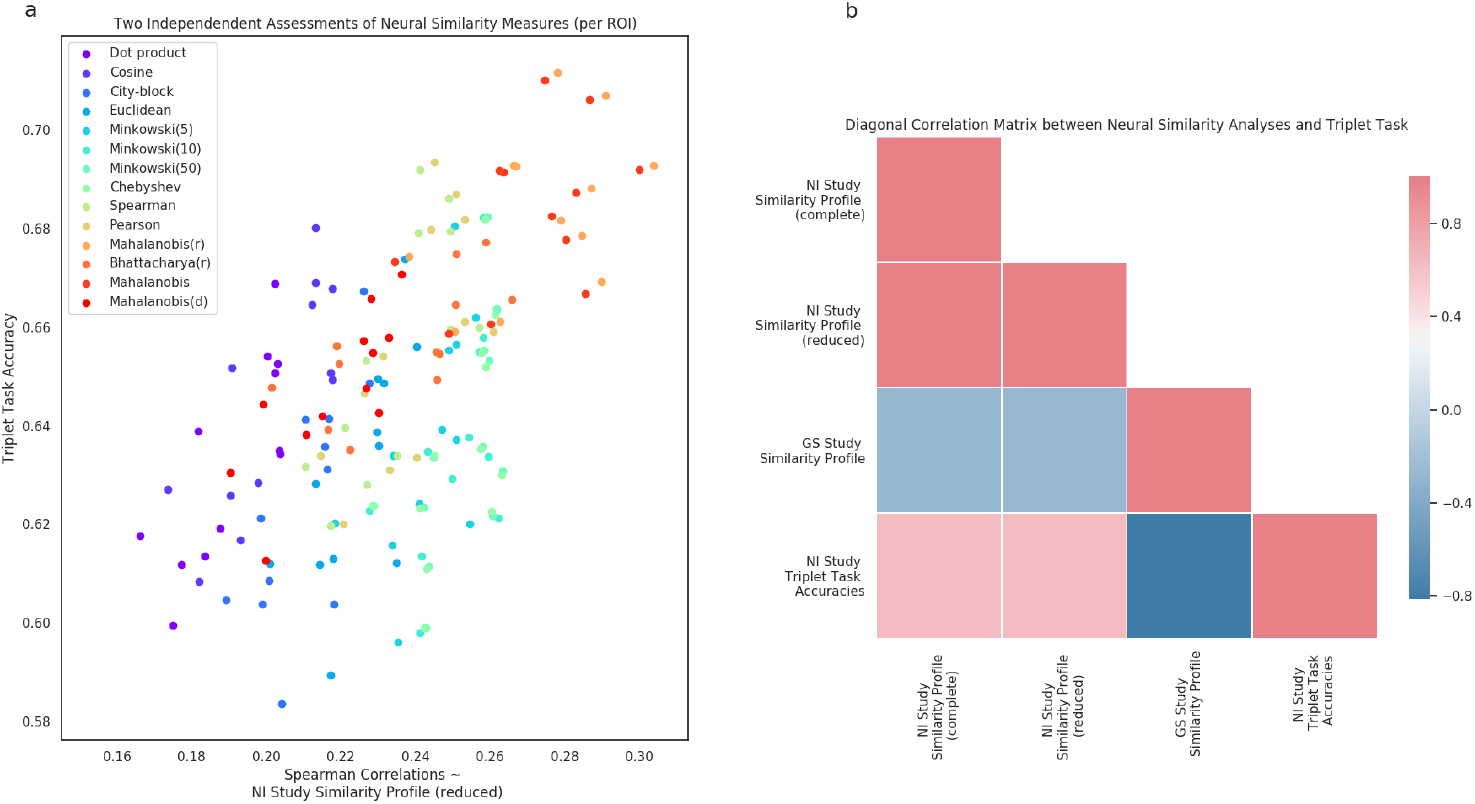
Triplet task accuracies correlate with NI Study similarity profiles. In (a) each data point represents one similarity measure per region of interest. The Spearman correlations in (a) have been recalculated with the removal of held-out pairs used in the triplet task (where each pair is the standard and the correct probe), thus termed NI Study similarity profile (reduced) (see Materials and Methods). In (b) we Pearson correlate the similarity profiles from the Neural Similarity analysis with the accuracies derived from the triplet task as well as with each other. NI Study similarity profile (complete) and GS Study similarity profile are the same Spearman correlations as displayed in Figure 3a (see Materials and Methods).

The triplet task allows a separate evaluation of the similarity measures of interest by comparing the accuracies in such a task to the similarity profile of the NI Study (Figure 5a); Pearson correlation of *r*(12) = 0.63, *p* = 0.017, across the fourteen similarity measures of interest. For this association, the scatterplot in Figure 5a shows the variance associated to the twelve regions of interest presented above. Measures like Mahalanobis and Mahalanobis(r) clearly do best; in line with the original similarity profile of the NI Study reported in the Neural Similarity analysis (Figure 3a). The similarity profile correlations were adjusted to account for the held-out pairs from the triplet task (with standard and correct probe removed), thus termed *(reduced)* in contrast to the original profile and reported here as *(complete)* (see Materials and Methods). In Figure 5b, all the similarity profiles are related amongst each other and with the triplet task accuracies. Most notably, the bottom row of the diagonal matrix displays how the triplet task accuracies also Pearson correlate negatively with the GS Study Similarity profile as in Figure 3d, *r*(12) = —0.81, *p* < 0.001. For comparison purposes, we also present the Pearson correlation of the triplet task accuracies with NI Study Similarity profile (complete), r(12) = 0.63,p = 0.016. The triplet task is thus an independent assessment of the validity of our neural similarity analysis.

These results clearly demonstrate that there is no circularity in our method of selecting similarity measures based on a decoding approach that approximates the information available in a brain state. In the triplet task, similarity measures that performed best in our neural similarity analysis also performed best in this novel task involving untrained classes.

More basic evidence against circularity claims is also presented in the SI; the best-performing classifier is a linear SVM for both the GS and NI study whereas we find dramatic differences in similarity profiles between studies. Clearly, similarity is not a simple recapitalization of classification.

## 3. Discussion

One fundamental question for neuroscience is what makes two brain states similar. This question is so basic that in some ways it has been overlooked or sidestepped by assuming that Pearson correlation captures neural similarity. Here, we made an initial effort to evaluate empirically which of several competing similarity measures is the best description of neural similarity.

Our basic approach was to characterize the question as a model selection problem in which each similarity measure is a competing model. The various similarity measures (i.e., models) competed to best account for the data, which was the confusion matrix from a classifier (i.e., decoder) that approximated the information present in a brain region of interest. The motivation for this approach is that more similar items (e.g., a sparrow and a robin) should be more confusable than dissimilar items (e.g., a sparrow and a moped). Thus, the test of a similarity measure, which is a pairwise operator on two neural representations, is how well its predicted neural similarities agree with the classifier’s confusion matrix.

At this early juncture, basic questions, such as whether different brain regions use different measures of similarity and whether the nature of neural similarity is constant across studies remained unanswered. Our results indicated that the neural similarity profile (i.e., the pattern of performance across candidate similarity measures) was constant across brain regions within a study, though strongly differed across the two studies we considered. Furthermore, Pearson correlation, the *de facto* standard for neural similarity, was bested by competing similarity measures in both studies.

Support for the validity of our method came from the follow-on triplet task in which we tested the ability of the similarity measures to select which of two probe items was most neurally similar to a comparison item. Similarity measures that performed best at this task (by selecting the probe that matched the comparison in stimulus shape) were those that also performed best under our decoding approach to evaluating neural similarity, despite the fact that the stimuli and classes used in the triplet were withheld from the decoding analyses. These results establish that our method of evaluating similarity measures selects measures that generalize well to novel tasks and stimulus classes. It also highlights that similarity and classification are distinct functions.

Accordingly, we report results in the SI in which the best performing similarity measures vary while the best performing classifier remains con stant, providing an illustration of how similarity and classifier performance can diverge. Of course, despite similarity and classification being distinct, the classifier used to estimate the information present in a brain region could bias the results. We recommend the procedure we followed: Consider a variety of classifiers and choose the best performing classifier independently of how the neural similarity measures perform (see SI). In practice, this means that an advance in classifier techniques would invite reconsidering how neural similarity measures perform.

One question is why the neural similarity profile would differ across studies. There are host of possibilities. One is that the nature of stimuli drove the differences. The stimuli in the GS study were designed to be psychologically separable, consisting of four independent binary dimensions (color: red or green, shape: circle or triangle, size: large or small, and position: right or left). These stimuli were designed to conform to a Euclidean space so that cognitive models assuming such similarity spaces could be fit to the behavioural data. Accordingly, in our analyses, the neural similarity measures from the Minkowski family (including Euclidean) performed best. In contrast, the NI study consisted of naturalistic stimuli (photographs) that covaried in a manner not easily decomposable into a small set of shared features. One possibility is that these types of complex feature distributions are better paired with the Mahalanobis measure (cf. [48]). Of course, task also varied with stimuli which offers yet another possible higher-level explanation for the differences observed in neural similarity performance. For example, the task in the GS study emphasized analytically decomposing stimuli into separable dimensions whereas holistic processing of differences was a viable strategy in the NI study. In general, different tasks will require neural representations that differ in their dimensionality or complexity [26], which has ramifications for what similarity measure is most suitable.

A host of other concerns related to data quality may also influence how similarity measures perform. The nature of fMRI BOLD response itself places strong constraints on the types of models that can succeed [49], which suggests that future work should apply the techniques presented here to other measures of neural activity. Regardless of the measure of neural activity, more complex models of neural similarity will require higher quality data to be properly estimated. For example, measures such as Mahalanobis or Bhattacharyya need to estimate inverse covariance matrices. These matrices grow with the square of the number of vector components which approaches both numerical and statistical unreliability when the number of components approaches the number of observations. For these reasons, we optimized the number of top features (i.e., voxels) separately for each similarity measure (see Materials and Methods), except in the searchlight analysis where this was not possible. We also considered regularized versions of similarity measures, such as Mahalanobis(d), that should be more competitive when data quality is limited.

Although the similarity measures considered are relatively simple, they make a host of assumptions that are theoretically and practically consequential. For example, angle measures, such as Pearson correlation, are unconcerned with differences in the overall level of neural activity, an assumption that strongly contrasts with magnitude measures, such as those in the Minkowski family (e.g., Euclidean measure). Therefore, the choice of similarity measure is central to any mechanistic theory of brain function and has practical ramifications when analyzing neural data, such as when characterizing neural representation spaces. In this light, operations that may seem routine, such as normalizing data in various ways, can affect the interpretation of results. For example, vector cosine only differs from dot product by virtue of normalizing by the magnitude of the two state vectors.

As mentioned previously, the space of possible similarity measures is un-countably infinite and new measures routinely enter the literature [50, 46]. In line with our main results, sometimes new measures like crossnobis perform well, and sometimes they fail [51]. Here, we aimed to include representative measures from the main families of similarity measures we identified (see Figure 1, left side). Others are free to replicate our analyses with alternative sets of measures.

Although we focus on the BOLD response, our approach applies equally to other neural measures, such as single-unit recordings. One important open question is whether the same similarity measures perform well across measures that differ dramatically in terms of spatial and temporal resolution, as well as the aspects of neural activity they capture. Likewise, our approach can be applied to complex artificial neural networks, such as deep convolutions neural networks (CNN), which have become popular in neuroscience by virtue of their ability to track neural activity along the ventral stream during object recognition tasks [52]. In standard neural networks, the basic mathematics of integrate-and-fire artificial neurons (i.e., units) can be viewed as a similarity operation, namely a dot product between the weight representation of the unit and the activity pattern at the previous layer. Alternatively, many of the other similarity functions we considered are differentiable and could be used in CNNs trained through backpropagation to perhaps provide better performance and agreement with neural measures. The question of which similarity functions manifest at the unit level of a CNN vs. at a larger organizational level recapitulate the previous discussion of the human brain.

In conclusion, we took a step toward determining what makes two brain states similar. Working with two fMRI datasets, we found that the best performing similarity measures are common across brain regions within a study, but vary across studies. Furthermore, we found that the *de facto* similarity measure, Pearson correlation, was bested in both studies. Although follow-up work is needed, the current findings and technique suggest a host of productive questions and have practical ramifications, such as determining the appropriate measure of similarity before conducting a neural representational analysis. In time, efforts making use of this and similar approaches may lead to mechanistic theories that bridge neural circuits, related measurement data, and higher-level descriptions.

## 4. Materials and Methods

### 4.1. Datasets

The analyses are based on two previous fMRI studies: a study that presented simple geometric shapes (GS) to participants [28] and a study that presented natural images (NI) to participants [6]. The GS study consisted of a visual categorization task with 20 participants and the NI study of a 1-back size judgment task with 14 participants. Descriptions of the tasks and acquisition parameters can be consulted in the SI. For further information, the reader should consult the source citation directly.

### 4.2. Classification analysis

Pattern classification analyses were implemented using PyMVPA [53], Scikit-Learn [54], and custom Python code. The input to the classifiers were least squares separate (LS-S) beta coefficients for each presentation of a stimulus [55] (see SI). Three classifiers were used for the pattern classification: Gaussian naïve Bayes, *k*-nearest neighbor, and linear support vector machine (SVM). The output of one of these classifiers was to be chosen as the best representation of the underlying similarity matrix to which all other similarity measures would be compared to (see Neural similarity analysis below). The linear SVM was implemented with the *Nu* parametrization [56]. This *Nu* parameter controls the fraction of data points inside the soft margin; the default value of 0.5 was used for all classifications. The *k*-nearest neighbor classifier was implemented using five neighbors. No hyperparameters required setting for the Gaussian naïve Bayes classifier.

To pick the best-performing classifier, classification was conducted on the whole-brain (no parcellation into distinct ROIs) for each study independently. All classifiers were trained with leave-one-out *k*-fold cross-validation, where *k* was equal to the number of functional runs for each participant in each study (e.g. six runs in the GS study or sixteen runs in the NI study). To do feature selection on voxels, all voxels were ordered according to their *F* values computed from an ANOVA across all class (stimuli) labels. The top 300 voxels with the highest *F* values were retained based on classifier performance (i.e., accuracy) on the test run. For these classifiers, accuracy was computed across all classes (16 classes for the GS study and 54 classes for the NI study) with a majority vote rule across all computed decision boundaries (for classifiers where this is applicable like linear SVM). This means that random classification is equal to 6.25% for the GS study and 1.85% for the NI study for this whole-brain analysis. However, for all other classification analyses, accuracy is computed as mean pairwise accuracy across all classes, which means that random classification is equal to 50%. The best-performing classifier was selected as the classifier with highest mean accuracy (mean across participants) in the GS and NI study, independently. Classifier accuracies (i.e., confusion matrices) were multiplied by negative one for the neural similarity analysis explained. This was done so that they would correlate positively with the similarity measures and facilitate presentation of results.

The following analysis was performed for each of the 110 ROIs that are described in the SI. To train the classifiers leave-one-out *k*-fold cross-validation was also used. Within each fold, a (randomly) picked validation run was used to tune the number of features (i.e., voxels) that would be selected for that fold. Thus, feature selection was done within each fold. To do this feature selection, all voxels were ordered according to their *F* values computed from an ANOVA across all class (stimuli) labels. This step aids classifier performance because it preselects task relevant voxels (as opposed to item discriminative voxels). It is important to note that these ANOVAs were computed on the training runs but not on the validation run nor on the held-out test run, to avoid overfitting. The top *n* voxels with the highest *F* values were retained based on classifier performance (i.e., accuracy) on the validation run. Scipy’s *minimizescalar* function [57] was used to optimize this validation run accuracy with respect to the top *n* voxels. After picking the top *n* voxels, the classifiers were trained on both the training runs and the validation run. Subsequently, the classifiers were tested on the held-out test run for that fold. This classification analysis was done for all possible pairwise classifications for each study (i.e., 120 pairwise classifications in the GS study and 1431 pairwise classifications in the NI study). From this analysis, the pairwise classification accuracies were retained for both the validation run and the test run for each fold. Further ROI selection (top twelve ROIs reported in the Results) is described in the SI.

### 4.3. Neural similarity analysis

The goal of this analysis was to compare the representation of different similarity measures in the brain. The regions considered here are the ones reported in the Results and described in the secondary ROI selection section in the SI. The comparison criterion was chosen as Spearman correlation between all pairwise similarities and the classification accuracies mentioned above. This criterion was used since it avoids scaling issues. To achieve this, first all pairwise similarities (i.e., for all pairs of stimuli) were computed from the training runs defined in the classification analysis not including the validation run. Incidentally, feature selection was also realized here. In the same fashion as in the classification analysis, all voxels were ordered according to their *F* values computed from an ANOVA across all class (stimuli) labels. Then, the top *n* voxels with the highest *F* values were retained based on Spearman correlation of the similarities with the validation run accuracies of the classifier that were previously computed. After picking the top *n* voxels, the similarities were computed across training runs and validation run for those voxels. These similarities were then used to compute the final Spearman correlation with the classifier test run accuracies. Conducting feature selection for the similarity measures is important because different measures leverage information differently.

This analysis parallels the classification analysis in every way except that instead of optimizing model accuracy, here the optimization criterion was model correlation (i.e., Spearman correlation) with the previously computed pairwise classifier accuracies.

### 4.4. Mixed effects models

A mixed effects model was performed with the lme4 package [58] for each study with Spearman correlations from the neural similarity analysis (i.e., similarity profile) as the response variable. The models contained fixed effects of similarity measure, linear SVM accuracy, participant, and ROI. Linear SVM accuracy, participant, and ROI variables only serve to account for variance and obtain better estimates. The models also contained random effects of ROI (varying per participant) and of similarity measure (varying per ROI). Model comparisons were performed between the full model and a null model without any similarity measures.^2^

### 4.5. Post hoc searchlight analysis

Searchlight analyses [59] have become an increasingly popular multivariate tool for spatial localizations of brain activations in recent years. This analysis is based on the definition of a sphere with radius in millimeters (or cube with radius in number of voxels) that computes a statistic, centered on each voxel of interest, using as input only the voxel values that fall within the confines of the predefined sphere. Depending on the number of voxels considered, this analysis can be computationally expensive. Thus for reasons of computational tractability, a searchlight analysis was not used as the primary analysis but as a *post hoc* tool to inquire over the spatial specificity of certain measures of interest commonly used in the literature such as Euclidean, Pearson correlation and Mahalanobis [16]. Since optimizing the searchlight radius for each voxel is not feasible with current computational resources – to equate measure complexity by feature selection as done in the main analysis – the searchlight radius was set to 3 voxels. The analysis was done only for Euclidean, Pearson correlation, and Mahalanobis(r). This searchlight analysis was done within the union of the top 10 ROIs across both studies (see Secondary ROI selection above) in the native space of each subject using PyMVPA’s searchlight function. For each voxel, the similarity matrices were Spearman correlated with the best performing classifier in the same fashion as in the main analysis above. For each study, the statistical maps of Euclidean and Mahalanobis(r) were compared to the statistical map of Pearson correlation, using it as a baseline measure. All maps were transformed to MNI space for this comparison. The threshold-free enchancement (TFCE) corrected *p* values for the paired *t* statistics were computed with FSL’s randomise function with 5000 permutations. Only *t* statistics that presented TFCE corrected *p* values below 0.001 were considered as significant. This more conservative threshold was based upon this being a *post hoc* analysis (i.e., supposing all 17 measures would have been compared against Pearson correlation, then the appropriate Bonferroni corrected threshold would have been *p* = 0.05/17 ≈ 0.0029).

### 4.6. Triplet task

In this task, a stimulus is chosen as a standard and paired with a correct probe. These pairs of standard and correct probe are designated as held-out pairs for reasons that will seem obvious below. The correct probes are chosen so as to share a common dimension with the standard. For the NI study, this was possible since shape (or silhouette) and category were two orthogonal dimensions that were part of the experimental setup of the fifty-four stimuli in the original study. In that study, six categories of stimuli were orthogonal to nine types of shape or silhouette (see SI). Subsequently, incorrect probes were chosen on the basis of not sharing values for either the shape or category dimensions for both the standard and correct probe. Thus, if basing the choice of correct probe on agreement with the shape dimension, then thirty-two (out of fifty-four) stimuli remain as choices for incorrect probes (i.e., shape triplet task). However, if the choice of correct probe is agreement with category, then thirty-five (out of fifty-four) stimuli remain as choices for incorrect probes (i.e., category triplet task). Either way, the task then consists of comparing the similarity between standard and correct probe with similarity between standard and incorrect probe for a given similarity measure. If the similarity measure is higher between standard and correct probe than it is for standard and incorrect probe, then the outcome of such a comparison is labelled with value one, otherwise zero. This operation was done for all possible incorrect probes and accuracy was computed as the number of outcomes equal to one divided by the number of incorrect probes (thirty-two for the shape triplet task and thirty-five for the category triplet task). This procedure was repeated for all possible pairs of standard and correct probe per similarity measure (out of 14 similarity measures reported in the Results), per run, per region of interest (out of the 12 ROIs reported in the Results), and per subject.

Accuracies computed for the triplet task should show performance of similarity measures that are in agreement with the similarity profiles from the neural similarity analysis. To adequately assess such an agreement, we recalculated the similarity profiles based on a subset of the original Spearman correlations used in the neural similarity analysis. The subset consisted of removing correlations for held-out pairs (standard and correct probe) that were being assessed in the triplet task. Such a procedure is necessary to claim independence between the neural similarity analysis and the triplet task and avoid inflated correlations between tasks. This subset of Spearman correlations was used to calculate the similarity profile referred to as *NI Study Similarity Profile (reduced)*, whereas the original similarity profile was referred to as *NI Study Similarity Profile (complete)* in the Triplet Task subsection of the Result section.

## Data and code availability

For open access to the data or code please visit

1) Raw fMRI data for the GS Study: https://osf.io/62rgs/
2) Raw fMRI data for the NI Study: https://osf.io/qp54f/
3) Data and code for the neural similarity analysis: https://osf.io/5a6bd/

## Acknowledgements

From the Love Lab, we thank Olivia Guest, Kurt Braunlich, Brett Roads, Rob Mok, Adam Hornsby, and Xiaoliang Luo for their comments. We thank the authors of the NI and GS Studies for making their data publicly available. This research was supported by a scholarship from Consejo Nacional de Ciencia y Tecnologa (CONACYT) and an enrichment year stipend from The Alan Turing Institute to SBS and by the NIH Grant 1P01HD080679, Lever-hulme Trust grant RPG-2014-075, and Wellcome Trust Senior Investigator Award WT106931MA to BCL.

## Author contributions

BCL developed the study concept. BCL and SBS contributed to the study design. SBS performed the data analysis and interpretation under the supervision of BCL. AM and AP performed confirmatory checks of the results and auxiliary analyses. SBS drafted the manuscript. BCL and CA provided critical revisions. All authors approved the final version of the manuscript for submission.

## Competing financial interests

The authors declare no competing financial interests.

## Supplementary Information

### A. Task descriptions and fMRI parameters

#### A.1 Geometric shapes (GS) study

The GS study presented sixteen objects in total, which varied on four different binary features: (color: red or green, shape: circle or triangle, size: large or small, and position: right or left). Participants in this study were trained to do a categorization task. They were first trained on five objects of one category and four of the other (nine objects total during training) with twenty repetitions of each object. During the anatomical scan, participants saw four more repetitions of the training items as a refresher. Then during the functional scanning phase, participants were asked to categorize the nine familiar objects they saw during the training phase and seven novel objects they had not seen before. Each trial during the functional scanning phase lasted 10 seconds; 3.5 seconds where one of the sixteen objects (nine training stimuli and seven novel transfer stimuli) was presented after which a fixation cross was presented for 6.5 seconds. No feedback was provided during this phase. Each stimulus was presented three times within a run across six runs resulting in each stimulus being presented a total of eighteen times during the functional scanning phase except for one participant who only participated in five runs of the scanning phase.

Whole-brain imaging data were acquired on a 3.0T GE Sigma MRI system (GE Medical Systems). Structural images were acquired using a T2-weighted flow-compensated spin-echo pulse sequence (TR=3s; TE=68ms, 256×256 matrix, 1×1mm in-plane resolution) with thirty-three 3-mm thick oblique axial slices (0.6mm gap), approximately 20 off the AC-PC line. Functional images were acquired with an echo planar imaging sequence using the same slice prescription as the structural images (TR=2s, TE=30.5ms, flip angle=73, 64×64 matrix, 3.75×3.75 in-plane resolution, bottom-up interleaved acquisition, 0.6mm gap). An additional high-resolution T1-weighted 3D SPGR structural volume (256×256×172 matrix, 1×1×1.3mm voxels) was acquired for registration and cortex parcellation.

#### A.2. Natural images (NI) study

The NI study presented fifty-four objects in total, which varied in two ways. The 54 stimulus items were conceived to either be organized by category (6 categories: minerals, animals, fruits/vegetables, music, sports, or tools) or by their silhouette (9 silhouettes) which cut orthogonally across the category distinction. Participants in this study were asked to perform a 1-back real-world size judgment task (i.e., to respond according to whether the object on the previous trial was larger or smaller than the current image on screen). Participants were scanned on two separate sessions (different days). Each session consisted of eight functional scanning runs resulting in sixteen runs total except for one participant for which four of the runs of the first session were lost due to scanning issues. Each one of the fifty-four objects were presented twice within each run in a randomized sequence. This resulted in each object being presented a total of thirty-two times (or twenty-four times for the participant that only had twelve runs). On each trial, each object was presented for 1.5 seconds after which a fixation cross was presented for 1.5 seconds. Each run started with a fixation cross for fourteen seconds and ended with a fixation cross for fourteen seconds. Thirty-six fixation trials lasting three seconds each were also randomly presented within each run.

Data collection was performed on a 3T Philips scanner with a 32-channel coil at the Department of Radiology of the University Hospitals Leuven. MRI volumes were collected using echo planar (EPI) T2*-weighted scans. Acquisition parameters were as follows: repetition time (TR) of 2 s, echo time (TE) of 30 ms, flip angle (FA) of 90, field of view (FoV) of 216 mm, and matrix size of 72×72. Each volume comprised 37 axial slices (covering the whole brain) with 3 mm thickness and no gap. The T1-weighted anatomical images were acquired with an MP-RAGE sequence, with 1×1×1 mm resolution.

#### A.3. fMRI preprocessing

The original raw (NIfTI formatted) files from both studies were preprocessed and analyzed using FSL 4.1 [1]. Functional images were realigned to the first volume in the time series to correct for motion, co-registered to the T2-weighted structural volume, high-pass filtered (128s), and detrended to remove linear trends within each run. All analyses were performed in the native space of each participant.

#### A.4 Trial-by-trial estimates

For both studies, after preprocessing the fMRI data with FSL, the method suggested by Mumford et al. [2] known as LS-S (least squares separate) beta estimation was used to get a coefficient estimate for each individual presentation of each object. This method consists of calculating a general linear model for each object presentation with only two regressors; one regressor representing the effect of interest (the object presentation in question) and another regressor representing all other object presentations within the respective run. This procedure was done for each run separately to preserve as much statistical independence as possible between runs. Such a step is necessary for doing the multivoxel pattern analysis. After successfully estimating the object presentation coefficients within each run, these were then concatenated into a single 4D NIfTI formatted file. Furthermore, all runs were subsequently aligned to the last run within each study (e.g. the sixth run in the GS study or the sixteenth run in the NI study). The runs were then concatenated into a single 4D NIfTI formatted file for each participant within each study.

### B. Regions of interest from the Harvard-Oxford atlas

#### B.1 Initial region of interest (ROI) selection

The Harvard-Oxford cortical and subcortical structural atlases provided with FSL [1] were used to parcellate the different anatomical regions for each participant. A total of 110 regions of interest were used as masks that would be used in the multivoxel pattern analyses. The goal was to evaluate classifier accuracy across the whole brain (except for areas like cerebral white matter or the lateral ventricles). More areas could have been excluded based on a priori hypotheses of where similarity signals would arise. However, including areas where no signal was expected served as an informal control for the method and still retained the possibility that similarity signals could have been found in otherwise unexpected brain regions. The masks were transformed from MNI space to each participants native space. This masking by anatomical region can be considered the first part of a feature selection procedure. Feature selection was also done within each region of interest for each participant (see Materials and Methods). All regions from the Harvard-Oxford atlas were included in the analyses except for cerebral white matter, the lateral ventricles, left and right cerebral cortex, and the brain stem. This results in 48 cortical regions and 7 subcortical regions; doubling for lateralization results in the 110 regions of interest.

#### B.2 Cortical regions of interest

Frontal Pole, Insular Cortex, Superior Frontal Gyrus, Middle Frontal Gyrus, Inferior Frontal Gyrus (pars triangularis), Inferior Frontal Gyrus (pars opercularis), Precentral Gyrus, Temporal Pole, Superior Temporal Gyrus (anterior division), Superior Temporal Gyrus (posterior division), Middle Temporal Gyrus (anterior division), Middle Temporal Gyrus (posterior division), Middle Temporal Gyrus (temporooccipital part), Inferior Temporal Gyrus (anterior division), Inferior Temporal Gyrus (posterior division), Inferior Temporal Gyrus (temporooccipital part), Postcentral Gyrus, Superior Parietal Lobule, Supramarginal Gyrus (anterior division), Supramarginal Gyrus (posterior division), Angular Gyrus, Lateral Occipital Cortex (superior division), Lateral Occipital Cortex (inferior division), Intracalcarine Cortex, Frontal Medial Cortex, Juxtapositional Lobule Cortex (formerly Supplementary Motor Cortex), Subcallosal Cortex, Paracingulate Gyrus, Cingulate Gyrus (anterior division), Cingulate Gyrus (posterior division), Precuneous Cortex, Cuneal Cortex, Frontal Orbital Cortex, Parahippocampal Gyrus (anterior division), Parahippocampal Gyrus (posterior division), Lingual Gyrus, Temporal Fusiform Cortex (anterior division), Temporal Fusiform Cortex (posterior division), Temporal Occipital Fusiform Cortex, Occipital Fusiform Gyrus, Frontal Operculum Cortex, Central Opercular Cortex, Parietal Operculum Cortex, Planum Polare, Heschl’s Gyrus (includes H1 and H2), Planum Temporale, Supracalcarine Cortex, & Occipital Pole.

#### B.3 Subcortical regions of interest

Thalamus, Caudate, Putamen, Pallidum, Hippocampus, Amygdala, & Accumbens.

#### B.4 Secondary ROI selection

The 110 ROIs were rank ordered by mean classifier accuracy (mean across participants) within each study. Subsequently, the union of the top ten ROIs was selected for the neural similarity analysis. This procedure was done to ensure that the ROIs used to evaluate the similarity measures was based on brain areas with adequate signal-to-noise ratio. The 12 ROIs as reported in the Results were left and right intracalcarine cortex (CALC), left and right lateral occipital cortex (LO) inferior and superior divisions, left and right lingual gyrus (LING), left and right occipital fusiform gyrus (OF), and left and right occipital pole (OP).

### C. Classifier selection

The best performing classifier was chosen out of three candidates; Gaussian naïve Bayes (GNB), *k*-nearest neighbor (KNN), and linear support vector machine (SVM). These classifiers were chosen because they are commonly used in data analysis, both inside and outside the field of neuroimaging, and they compute classification in very distinct ways (see [3]).

The linear SVM classifier was the clear winner across both studies, thus was chosen as our gold standard approximation to the brain’s similarity measure. The performance of the linear SVM classifier compared to the other two classifiers is shown in Table C1.

**Table C1.**
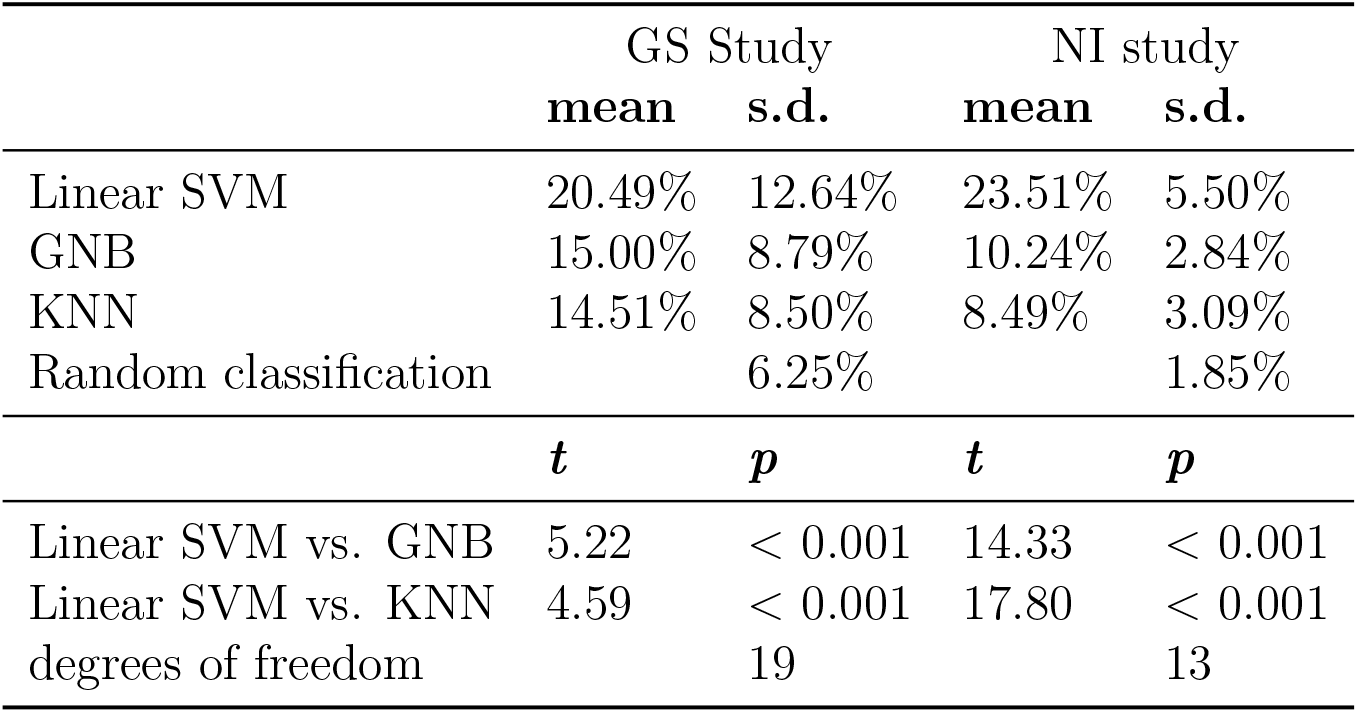
Linear SVM is best-performing classifier in both studies. Top panel shows mean accuracy and standard deviations (s.d.) (across participants) for each classifier. Bottom panel shows t-tests comparing the best-performing classifier (linear SVM) to the other two classifiers.

In addition to comparing the performance of the classifiers judged by their performance accuracy, the confusion matrices between classifiers – from the same analysis – were also compared. Although the classifiers are quite distinct algorithmically speaking, extreme differences between their confusion matrices would be unlikely. Indeed it was the case that the average correlations (averaged across subjects) were all significantly above zero for both studies. In the GS study, linear SVM correlated highest with GNB (m = 0.47, s.d. = 0.172, *t*(19) = 12.01, *p* < 0.001), second highest with KNN (m = 0.37, s.d. = 0.197, *t*(19) = 8.21, *p* < 0.001), and GNB correlated with KNN in third place (m = 0.32, s.d. = 0.195, *t*(19) = 7.06, *p* < 0.001). In the NI study, linear SVM correlated highest with GNB (m = 0.35, s.d. = 0.072, *t*(13) = 17.55, *p* < 0.001), second highest with KNN (m = 0.29, s.d. = 0.080, *t*(13) = 13.06, *p* < 0.001), and GNB correlated with KNN in third place (m = 0.22, s.d. = 0.091, *t*(13) = 8.94, *p* < 0.001). These results provide supplementary support for choosing linear SVM as the brain’s gold standard for these two datasets given that it’s confusion matrix correlates highest with the confusion matrices of the other two classifiers.

Thus, the linear SVM classifier was optimized for each of the initial 110 ROIs. The ROIs were rank-ordered in terms of accuracy in each study and the union of the top 10 ROIs across both studies was: left and right intracalcarine cortex (CALC), left and right lateral occipital cortex (LO) inferior division, left and right lateral occipital cortex (LO) superior division, left and right lingual gyrus (LING), left and right occipital fusiform gyrus (OF), and left and right occipital pole (OP). This resulted in a secondary ROI selection of 12 ROIs with best (linear SVM) classifier accuracy.

Classifications were performed pairwise for this analysis and thus random classification was expected at 50% for both studies (see Materials and Methods). The mean accuracy for the linear SVM classifier in the 12 regions of interest was 59.47% (s.d. = 7.97%) in the GS study and 78.43% (s.d. = 7.41%) in the NI study. The best-performing classifier (linear SVM) was performing above 50% chance level in both studies; *t*(19) = 5.18, *p* < 0.001, in the GS study and *t*(13) = 13.84, *p* < 0.001, in the NI study (degrees of freedom are based on number of participants for each study). This provides reassurance that the ROIs that were selected indeed have information regarding stimuli presentation. Classification accuracy for the NI study was higher than in the GS study *t*(32) = 6.82, *p* < 0.001, showing a potential difference in data quality due to the higher number of observations per stimuli in the NI study (see Materials and Methods).

### D. Similarity measures

The following similarity measures were evaluated: dot product, cosine distance, city-block (Manhattan), Euclidean, three variants of Minkowski (with norms 5, 10 and 50), Chebyshev, Spearman correlation, Pearson correlation, three variants of Mahalanobis, three variants of Bhattacharyya, variation of information, and distance correlation. City-block, Euclidean, Minkowski, Chebyshev, Mahalanobis, Bhattacharyya and variation of information are proper distance metrics; to convert them to similarity measures they were multiplied by minus one. Other linking functions between similarities and distances are possible, as in a negative exponential [4], but not relevant here since our optimization criterion was Spearman correlation. The three variants of Mahalanobis and Bhattacharyya were due to the way the sample covariance matrix was regularized; either no regularization, Ledoit-Wolf shrinkage (implemented through Scikit-Learn, [5, 6] or diagonal regularization. Diagonal regularization was defined as the sample covariance matrix with all the off-diagonal elements set to zero (see below); such as measure is also known as the normed Euclidean distance. Note that city-block, Euclidean, and Chebyshev are also special cases of the Minkowski measure where the norms are set to one, two and infinity, respectively. To keep calculations consistent across all similarity measures, vector representations for each stimulus were defined as the mean vectors across trial presentations for that stimulus. Below are the equations for each similarity measure and the covariance matrix regularization procedures.

In constructing the similarity profiles, we only used similarity measures that presented a mean Spearman correlation within three median absolute deviations away from the group average (group refers to measures here). Measures that did not meet these criteria were considered outliers (these measures were close to zero mean Spearman correlation). The median Spearman correlation across the 18 similarity measures evaluated was 0.203 for the GS study 0.125 and for the NI study and their median absolute deviation was 0.0482 for the GS study and 0.0234 for the NI study. The mean Spearman correlations (across participants) and the standard deviations for the measures that were more than three median absolute deviations away from the group average were: Bhattacharya without covariance matrix regularization (mean = 0.001 and s.d. = 0.004 for the GS study, mean = 0.0002 and s.d. = 0.0006 for the NI study), Bhattacharya (d) (with diagonal regularization) (mean = −0.0005 and s.d. = 0.003 for the GS study, mean = −0.0001 and s.d. = 0.0007 for the NI study), variance of information (mean = −0.04 and s.d. = 0.037 for the GS study, mean = −0.012 and s.d. = 0.004 for the NI study), and distance correlation (mean = −0.037 and s.d. = 0.026 for the GS study, mean = −0.0009 and s.d. = 0.0038 for the NI study). These statistics were computed across the 110 original ROIs.

Below is a list of the equations for each measure considered.

For two classes represented as vectors

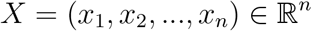

and

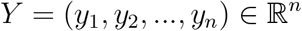

where each component is computed as the arithmetic mean across m observations (trial-by-trial β coefficients) per class, per run, and *n* is the number of voxels. This notation is valid except for where these vectors show subscripts denoting individual observations as opposed to mean vectors (this is only the case when discussing distance correlation).

#### Dot product

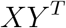

#### Cosine distance

The (negative) cosine distance is:

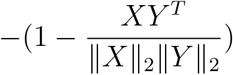

where || · ||_2_ denotes the L2 (Euclidean) norm.

#### Minkowski distance

The (negative) Minkowski distance is:

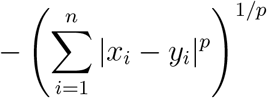

For the city-block distance *p* = 1, for the Euclidean distance *p* =2, and for the Chebyshev distance *p* = ∞.

#### Pearson correlation

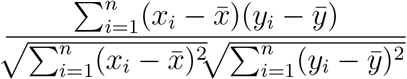

where 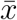 and 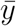 are the component-wise arithmetic means of vectors *X* and *Y*, respectively.

#### Spearman correlation

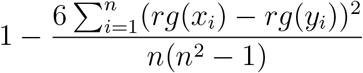

where *rg*(*x_i_*) and *rg*(*y_i_*) are the ranks of the values *x_i_* and *y_i_*, respectively. This formulation assumes distinct integer rankings.

#### Mahalanobis distance

The (negative) Mahalanobis measure between two random vectors coming from the same multivariate normal distribution is:

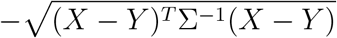

where Σ is the *n × n* covariance matrix between voxels.

#### Bhattacharyya distance

The (negative) Bhattacharyya measure between two multivariate normal distributions 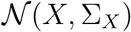 and 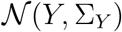, where each voxel covariance matrix Σ_*X*_ and Σ_*Y*_ is estimated separately for each class *X* and *Y*, respectively, is:

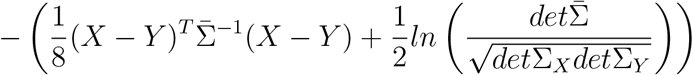

where

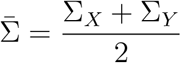

#### Distance correlation

The distance correlation is equal to 1 when *X* and *Y* span the same linear subspace under some linear transformation and 0 when *X* and *Y* are independent. It is defined as:

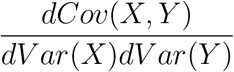

where *dCov*^2^(*X,Y*) is

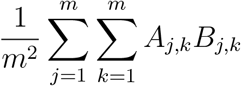

and *d Vav*^2^(*X*) is

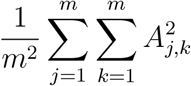

where *A_j,k_* is the matrix computed from doubly-centering the matrix *a_j,k_* (subtracting row and column means while adding the grand mean), where

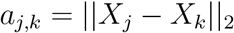

Thus, *B_j,k_* is computed from *b_j,k_*, where

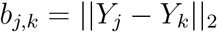

These pairwise distance matrices are computed from distances between observations.

#### Variation of information

For two classes *X* and *Y* represented as two multivariate Gaussian distributions, the (negative) Variation of information is

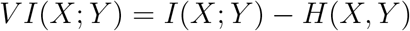

where *H*(*X*) is the entropy of *X* and *I*(*X; Y*) is the mutual information between *X* and *Y*.

For a multivariate Gaussian *X, H*(*X*) is:

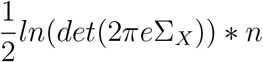

where *n* is the number of observations. The mutual information between *X* and *Y* is:

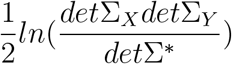

where Σ*

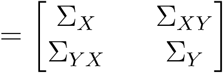

and Σ_*XY*_ is the between-class voxel covariance matrix. Σ_*YX*_ is the transpose of Σ_*XY*_.

#### Covariance matrix regularization

Two types of covariance matrix regularization were used for the Mahalanobis distance: diagonal regularization and Ledoit-Wolf regularization.

#### Diagonal regularization

Diagonal regularization for a covariance matrix Σ was computed as Σ o I, where o is the hadamard product (element-wise multiplication) and I is the identity matrix.

The distance measure that comes as a result of this type of regularization, when applied to the covariance matrix of the Mahalanobis distance, is also known as the normed Euclidean distance.

#### Ledoit-Wolf regularization

Ledoit-Wolf regularization for a covariance matrix Σ was computed as:

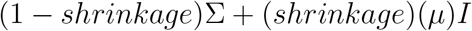

where *μ* = *trace*(Σ)/*n* and the optimal shrinkage parameter is a value between 0 and 1 estimated according to the derivation in [5].

### E. Post hoc searchlight analysis

Supplementary Figure 1 presents voxels where both the Euclidean measure and the Mahalanobis(r) measure outperformed Pearson correlation.

**Supplementary Figure 1:**
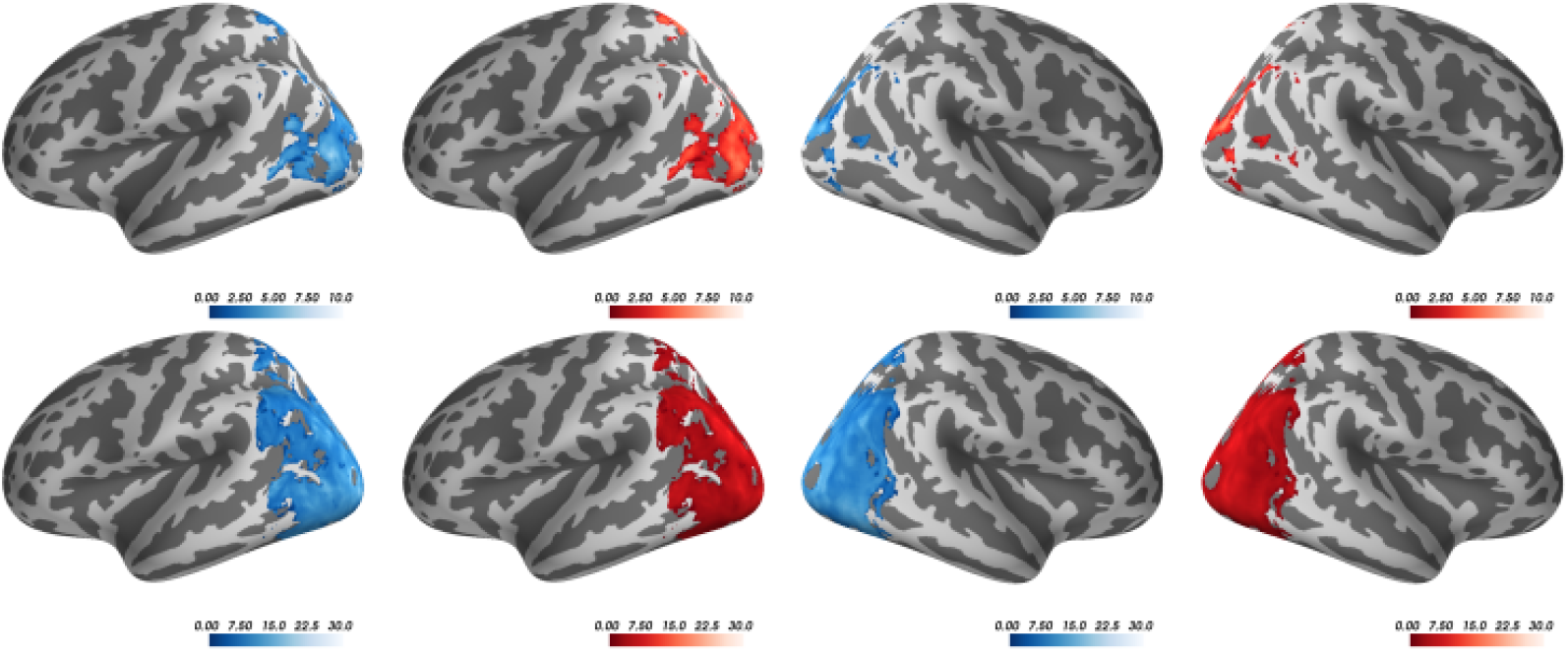
Voxels where Euclidean & Mahalanobis(r) overlap (outperforming Pearson). Lateral views of the left and right hemispheres for the GS study (top row) and the NI study (bottom row) displaying *t* statistics where both the Euclidean measure (blue) and the Mahalanobis(r) measure (red) outperformed the Pearson correlation measure. The *t* statistics were based on a searchlight analysis of Spearman correlations of each measure with each voxel’s SVM confusion matrix (see Materials and Methods). Only displaying *t* statistics where *p* < 0.001 for paired sample *t*-tests, TFCE corrected; computed with FSL’s randomise function with 5000 permutations, using as a mask the 12 ROIs with best accuracy (see Materials and Methods).

## Footnotes

1 Anisotropic measures should not be confused with asymmetric measures; the latter gives different values based on which stimulus is measured first [34, 7].

2 A full model that included both studies was not possible due to convergence issues.

